# Defective proinsulin handling modulates the MHC I bound peptidome and activates the inflammasome in β-cells

**DOI:** 10.1101/2021.12.20.472059

**Authors:** Muhammad Saad Khilji, Erika Pinheiro-Machado, Tina Dahlby, Ritchlynn Aranha, Søren Buus, Morten Nielsen, Justyna Klusek, Pouya Faridi, Anthony Wayne Purcell, Thomas Mandrup-Poulsen, Michal Tomasz Marzec

**Author notes:** **Corresponding Author** Michal Tomasz Marzec. (M.S.K); (T.M.-P); (M.S.K); (R.A); (A.W.P); (P.F), (M.S.K), (E.P.-M), (T.D), (S.B), (M.N), (J.K), (M.T.M).

## Abstract

**Background:** How immune-tolerance is lost to pancreatic β-cell peptides triggering autoimmune type 1 diabetes is enigmatic. We have shown that loss of the proinsulin ER chaperone glucose-regulated protein (GRP) 94 leads to mishandling of proinsulin, ER stress and activation of the inducible proteasome. We hypothesize that inadequate ER proinsulin folding capacity relative to biosynthetic need may lead to an altered β-cell MHC-I bound peptidome and inflammasome activation, sensitizing β-cells to immune attack.

**Methods:** We used INS-1E cells with or without GRP94 knockout (KO), or in the presence or absence of GRP94 inhibitor PU-WS13 (GRP94i, 20µM), or exposed to proinflammatory cytokines interleukin (IL)-1β or IFNγ (15 pg/ml and 10 ng/ml, respectively) for 24 hours. RT1.A (rat MHC I) expression was evaluated using flow cytometry. The total RT1.A-bound peptidome analysis was performed on cell lysates fractionated by reverse phase high performance liquid chromatography (RP-HPLC) followed by liquid chromatography coupled with tandem mass spectrometry (LC-MS/MS). NALP1, nuclear factor of kappa light polypeptide gene enhancer in B-cells inhibitor alpha (IκBα), and (pro) IL-1β expression and secretion were investigated by Western blotting.

**Results:** GRP94 KO increased RT1.A expression in β-cells as did cytokine exposure compared to relevant controls. Immunopeptidome analysis showed increased RT1.A-bound peptide repertoire in GRP94 KO/i cells as well as in the cells exposed to cytokines. The GRP94 KO/cytokine exposure groups showed partial overlap in their peptide repertoire. Notably, proinsulin-derived peptides diversity increased among the total RT1.A peptidome in GRP94 KO/i along with cytokines exposure. NALP1 expression was upregulated in GRP94 deficient cells along with decreased IκBα content while proIL-1β cellular levels declined, coupled with an increased secretion of mature IL-1β. Our results suggest that limiting β-cell proinsulin chaperoning enhances RT1.A expression, alters the MHC-I peptidome including proinsulin peptides and activates inflammatory pathways, suggesting that stress impeding proinsulin handling may sensitize β-cells to immune-attack.

## 1. Introduction

Type 1 diabetes (T1D) is an immune mediated destruction of β-cells as a consequence of a break in tolerance towards β-cell self-peptides. How this break in tolerance is attained still remains to be answered. Recent evidences point to the emergence of neoepitopes as a consequence of β-cell stress (Baker, et al. 2018; Gonzalez-Duque, et al. 2018). As a professional secretory cells, β-cells are challenged with physiological ER stress during the production and secretion of insulin (Marré, et al. 2016). In addition to inherent physiological stress, β-cells can be challenged further with environmental factors associated with the onset of T1D including but not restricted to, viral infection, diet, cross-reactivity to bovine albumin and gut microbiota composition (Atkinson, et al. 1994; Dedrick, et al. 2020; Holmberg, et al. 2007; Karjalainen, et al. 1992). All these factors lack uniformity and loss of tolerance could be related to β-cells unspecific secretory stress that exceeds β-cells inherent capacity. Proinsulin (PI) is the major β-cell translated product as well as a major autoantigen in human and rodent model of type 1 diabetes (Nakayama 2011; Zhang, et al. 2008). Recent advances in protein analytical technologies have expanded the list of neo-autoantigens in type 1 diabetes, especially those derived from (pro) insulin. These include normal insulin peptides, hybrid insulin peptides (Baker, et al. 2018; Baker, et al. 2019) as well as post-translational modifications (PTM) of insulin chains (Sidney, et al. 2018). It would suggest that nonconventional processing and degradation of insulin may contribute to triggering the loss of the peripheral immune tolerance. During its processing from pre-prohormone to mature insulin, PI requires the assistance of enzymes and chaperones in the ER such as protein disulfide isomerases (PDIs) and glucose regulated protein (GRP) -94 and -78 to facilitate disulfide bonds formation and to attain its native state (Ghiasi, et al. 2019; Liu, et al. 2010; Liu, et al. 2018; Scheuner, et al. 2005). Diminished activity of ER chaperones such as GRP94 results in proinsulin mishandling, provokes compensated ER stress in the form of PERK activation (Ghiasi, et al. 2019) and activates inducible proteasomes (Khilji, et al. 2020a), potentially setting the stage for inflammasome activation and neoepitope formation and presentation. Studies from non-obese diabetic (NOD) mouse showed an increased β-cell stress before the onset of clinical diabetes (Tersey, et al. 2012) as well as HLA hyper expression and presence of insulitis in human islets from a recent onset of T1D in patients (Coppieters, et al. 2012), suggesting β-cell stress precedes inflammatory events in the pancreas.

The incidence of T1D is increasing among children and adolescents (Patterson, et al. 2019). Although genetics account for roughly 50 % of total T1D development risk, its increased incidence in the recent past suggest an increased influence of environmental factors. One of the hypothesis gaining the support is the role of β-cells dysfunction at multiple levels in their own demise (Cnop, et al. 2012; Lamb, et al. 2008; Ludvigsson 2006; Marré, et al. 2015; Marre, et al. 2018; Rewers and Ludvigsson 2016; Sims, et al. 2019). Among these are believed to be those that affect (pro) insulin-folding resources and induce ER stress (e.g. growth, puberty, pregnancy, insulin resistance) (Sun, et al. 2015). It has been known that ER stress can induce the secretion of the ER resident heat shock protein (HSP) family of chaperones that can be taken up by antigen presenting cells (APC) (Heath and Carbone 2009; Steinman, et al. 1983) allowing cross presentation and activation of CD8^+^ T-cells (Norbury, et al. 2004). Cross presentation thereby bypasses the conventional APC-Th cell dependent immune activation, and sensitizes target cells, exposing neoepitope on MHC I to T-cell attack. GRP94 (an HSP90 family member) has been found free- or IgG-bound circulating in people with T1D (Pagetta, et al. 2003) and can efficiently cross-present chaperoned peptides to CD8^+^ T-cells (Murshid, et al. 2010). This suggests that activation of ER stress in response to environmental factors, followed by (or leading to) secretion of GRP94-PI complexes, may trigger the β-cell-directed immune response Recently we discovered that the β5i subunit of the int-proteasome is over-expressed in GRP94 deficient β-cells and its inhibition restores proinsulin levels (Khilji, et al. 2020a). Employing GRP94 KO/i and cytokines IL-1β/IFNγ exposure to mimic ER or inflammatory stress, we found that GRP94 KO and cytokines increased RT1.A expression in β-cells. Moreover, the total RT1.A bound peptide repertoire increased upon GRP94 KO/i/cytokines exposure of β-cells including peptides derived from (pro) insulin. GRP94 KO also increased NALP1 inflammasome expression, reduced cellular IκBα levels and reduced proIL-1β protein content coupled with an increased mature IL-1β secretion, indicating activation of a β-cell inflammatory response.

## 2. Methods

### 2.1. Procurement of cell lines and reagents

The mouse hybridoma cell line OX-18 for generating anti-RT1.A (anti-rat MHC I) antibody was purchased from CellBank, Australia. The rat insulinoma cell line (INS-1E) was a kind gift from Claes Wollheim (University Medical Center, Geneva, Switzerland). Protein G resin was purchased from Agarose Bead Biotechnologies (Cat# 4RRPG-100). All other chemicals/reagents of HPLC/MS grade were purchased from commercial vendors and are listed in the supplementary table 1.

### 2.2. Cell culture and pelleting

The unchallenged INS-1E and their derived GRP94 KO and control KO (cells targeted with lentiviral particles still expressing GRP94 similar to unchallenged INS-1E) cells were cultured and maintained between passage numbers 54 - 72 in RPMI-1640 as described previously (Khilji, et al. 2020a). To perform immunopeptidomics, cells from confluent flasks were detached using EDTA (10 mM), centrifuged at 4000 rpm to pellet the cells. The cell pellets of 5×10^8^ per replicate per group were snap frozen in liquid nitrogen and stored at -80 ^°^C until later use for immune-affinity capture of RT1.A. The hybridoma cell line OX-18 was cultured in roller bottles in RF5 medium (RPMI-1640 supplemented with 5 % fetal bovine serum (FBS), 1 % Penicillin-Streptomycin, 2 mM MEM non-essential amino acids, 100 mM HEPES, 2 mM L-glutamine and 50 µM β-mercaptoethanol). The supernatants were collected and antibody harvested using Profinia purification system (Bio-rad^®^).

### 2.3. Drugs treatment /cytokines exposure

To induce GRP94 functional loss or mimic inflammatory conditions, INS-1E cells were treated with either 20 µM GRP94 inhibitor (GRP94i) PU-WS13 (EMD Millipore Corp.^®^) or exposed to 15 pg/ml recombinant human interleukin-1β (IL-1β, Sino Biological Inc.^®^) or 10 ng/ml recombinant rat IFNγ (R&D^®^) for 24 hours prior to harvesting cells for flow cytometry or immunopeptidomics.

### 2.4. Flow cytometry

The unchallenged or GRP94 KO/control KO INS-1E cells cultured in the absence or presence of drug/cytokines at the dose and time mentioned above were washed with FACS buffer (2 % FBS in PBS) and incubated with OX-18 for 1 hour at 4^°^ C. The cells were again washed with FACS buffer followed by incubation with phycoerythrin (PE)-labelled anti-mouse antibody for 30 minutes. The cells were then re-suspended in 300 µl FACS buffer for flow cytometry. The flow cytometry data for RT1.A was acquired by LSRII (BD Biosciences^®^). The data were analyzed using flowjo software (Tree Star Inc^®^). The cells were gated on SSC-A × FSC-A and FSC-H × FSC-A to exclude doublets and debris, and the mean fluorescence intensity (MFI) for each group was recorded at 585 nm.

### 2.5. Isolation of RT1.A bound complexes

The detailed procedures for antibody cross-linking, purification of MHC bound complexes and fractionation have been described previously (Purcell, et al. 2019). Briefly, the frozen INS-1E cell pellets were grinded using cryogenic mill (Retsch Mixer Mill MM 400) and lysed in buffer containing 0.5 % IGEPAL (Sigma-Aldrich^®^), 50 mM Tris pH 8, 150 mM NaCl and supplemented with protease inhibitor cocktail (Roche^®^). The lysate was ultra-centrifuged (Beckman Coulter Optima L90K 77LR271) at 40,000 x g, 4^°^C for 2 hours. The lysate supernatant was collected, and RT1.A complexes were immunoaffinity purified by passing through a column containing cross-linked OX-18 antibody coupled to protein G agarose resin (ABT Beads^®^). The bound complexes were washed with buffers with decreasing detergent and salt concentrations as detailed in (Purcell, et al. 2019). The peptides were eluted in 10 % acetic acid for fractionation.

### 2.6. Fractionation by reverse phase high performance liquid chromatography (RP-HPLC)

The eluents were fractionated through RP-HPLC on a C18 column (Chromolith Speed Rod, Merck-Millipore^®^) using ÄKTA micro HPLC system (GE Healthcare^®^) running a mobile phase consisting of buffer A (0.1 % trifluoroacetic acid (TFA)) and buffer B (80 % acetonitrile/0.1 % TFA). Flowrate was adjusted at 2 ml/min to collect peptides in 1 ml fractions in low-protein binding Eppendorf tubes (Eppendorf LoBind tubes, cat. no. EPPE0030108.116). The bound complexes were separated from β2 microglobulin and heavy chain using increasing concentration of buffer B. Peptide containing fractions were collected, vacuum concentrated on speedVac (Labconco^®^) to ∼5 µl and combined into 5 pools per replicate. The pools were reconstituted in 0.1% formic acid + 2% acetonitrile (final volume = 10 µl) and stored in -80 ^°^C until run on LC-MS/MS.

### 2.7. Mass spectrometry data acquisition and analysis

The pooled samples were sonicated, centrifuged at 15,000 RPM for 30 min and transferred to mass spectrometry vials, each pool spiked with Indexed retention time (iRT) peptides to adjust retention time. The samples were run on LC-MS/MS (Orbitrap fusion Tribrid, Thermo^®^) on data-dependent acquisition (DDA) mode using the settings described in (Faridi, et al. 2020).

The .raw files acquired from the system were run on PEAKS (PEAKS 8.5, Bioinformatics solutions Inc. ^®^) against rat proteome (Uniprot 8,133 reviewed entries by 30 November 2021). The parameters included error tolerance of 10 ppm and 0.02 Da tolerance for fragment ions, enzyme was set to none and the PTMs included oxidation, cysteinylation and deamidation (NQ). The list of peptides and proteins were exported as .csv files at 5 % FDR.

### 2.8. Bioinformatics analyses

The binding motifs were created using IceLogo (Maddelein, et al. 2015), Venn diagrams at BioVenn.nl (Hulsen, et al. 2008) and unsupervised clustering for alignment of RT1.A-bound 8-15 mer peptide sequences using Gibbscluster (Andreatta, et al. 2017). The binding motif is presented for two clusters due to limited peptide yield in some groups. For analysis of the two GRP94 KO clones, the duplicates were combined into quadruplets to avoid clonal variations. However, we focus on the peptides that are detected in both GRP94 KO clones at least in one out of two biological replicates to minimize clonal variation.

To construct artificial neural networks based model to predict receptor ligand interaction, the tool NNAlign (Nielsen and Andreatta 2017) was trained on combination of random and INS-1E peptides (excluding proinsulin partite) with maximum overlap for common motif changed to ‘8’. The proinsulin partite and random natural peptides were then run against the trained model and compared to get percentile rank values against the trained model.

### 2.9. Tryptic digestion of heavy chain

To confirm specificity of OX18 for both class Ia and class Ib associated peptides, we performed tryptic digestion of the heavy chain fragment obtained from RP-HPLC fractionation. Briefly, the heavy chain fragments were pooled, vacuum dried to reduce volume and digested overnight with 2 µg trypsin in S-trap columns. For peptide recovery, 50 mM tetraethyl ammonium bicarbonate buffer was added to the digest followed by 0.2 % formic acid and finally 50 % acetonitrile with 0.2 % formic acid. The sample was centrifuged in between each step. The total sample volume was reduced to ∼5 µl in vacuum centrifuge and reconstituted in 0.1 % formic, 2 % acetonitrile solution (final volume = 10 µl) to run on LC-MS/MS (Triple-TOF 6600 system, SCIEX^®^).

For proteomics analysis of the tryptic digest of heavy chains, the settings included MS1 error 15 ppm MS2 error 0.1 Da and PTMs included: fixed carbamidomethylation of cysteine variable: M(O), N-terminal acetylation. Data was exported for analysis at 5 % FDR as .csv files.

### 2.10. Western blotting

The Western blotting was performed following the procedure detailed in (Khilji, et al. 2020b). In brief, 4 million of the confirmed CRISPR/Cas-9 targeted GRP94 KO INS-1E and control KO cells (CRISPR/Cas-9 targeted INS-1E cells still expressing GRP94 similar to unchallenged INS-1E) were grown for 48 hours in T25 flasks. Before lysing cells, the supernatant was collected to estimate secreted IL-1β. The cells were then lysed in buffer supplemented with protease inhibitor cocktail (Life Technologies, Nærum, Denmark). The protein concentration was calculated using Bradford assay (Bio-Rad, Copenhagen, Denmark). Thirty µg of total protein was loaded for each sample on 4-12 % bis-tris gel and afterwards transferred to PVDF membrane (iBLOT2 system^®^). The membranes were blocked in 5 % milk in TBST buffer, incubated overnight with antibodies against proinsulin, tubulin, NALP1, IκBα, IL-1β and GRP94 as listed in supplementary table 1 along with the dilution used, followed by 1 hour incubation with anti-rat, anti-mouse or anti-rabbit antibodies depending on the primary antibody host. The membranes were developed after washing 3× in TBST. The chemiluminescent images were acquired using Azure^®^ Saphire Biomolecular Imager. For supernatant Western blots to estimate secreted IL-1β, the procedure was performed as described previously by (Ghiasi, et al. 2019). Briefly, 500 µl of media was centrifuged at 14000 rpm for 30 minutes using 10 kDa filter to concentrate IL-1β into ∼40 µl volume. The supernatants were then mixed with lysis buffer and loading buffer and run on gel as described above. The images were quantified for protein expression using ImageJ (v 1.52a) and the ratio for protein of interest to tubulin was used for statistical comparison. For IL-1β supernatants, only bands were quantified for statistical comparison.

### 2.11. Statistical analysis

The flow cytometry data was analyzed using paired t-test for treatment versus relevant control groups using statistical software GraphPad prism version 9.1.0 (La Jolla). For Western blots, unpaired t-test was used on quantified blots for statistical comparison.

### 2.12. Data availability

The complete data sets of peptides and sources proteins are uploaded on Electronic Research Data Archive at University of Copenhagen (KU/UCPH) and can be acquired by sending email to.

## 3. Results

### 3.1. INS-1E cells express both class Ia and class Ib region of RT1 system

Rat RT1 system has two MHC I regions. RT1.A contains the centromeric, classical Ia region whereas the telomeric, nonclassical Ib region is referred as RT1.C/E/M. OX-18 is known to cross react with non-classical Ib products (Jameson, et al. 1992), though class Ib products are generally poorly expressed in different tissues (Reizis, et al. 1997). We performed tryptic digestion of the heavy chain fragments from the fractionated eluent to explore which class I regions are captured by OX-18 antibody in INS-1E cells. We found classical Ia and non-classical Ib derived peptides in the tryptic digest suggesting both class Ia and Ib are expressed in INS-1E cells and captured by OX-18. However, we can’t differentiate the expression levels of individual RT1 regions in our experimental settings (supplementary table 2).

### 3.2. GRP94 KO or cytokines exposure increase RT1.A expression

To estimate if GRP94 KO or cytokines exposure altered overall RT1.A expression, INS-1E cells were cultured with/without cytokines IL-1β/IFNγ or in the presence of GRP94i or GRP94 KO. The cells under tested conditions were incubated with OX-18 antibody followed by incubation with PE-labelled secondary antibody. Unchallenged INS-1E cells showed considerable basal RT1.A expression with a mean fluorescence intensity (MFI) around 1000. As expected, exposure to cytokine IL-1β or IFNγ significantly upregulated RT1.A expression, while GRP94i slightly upregulated RT1.A expression (figure 1-A & C). Interestingly, GRP94 KO increased RT1.A expression (figure 1-B & D) while GRP94i treatment of INS-1E cells showed a modest upregulation in MFI (*p-value* = 0.08).

**Figure 1.**
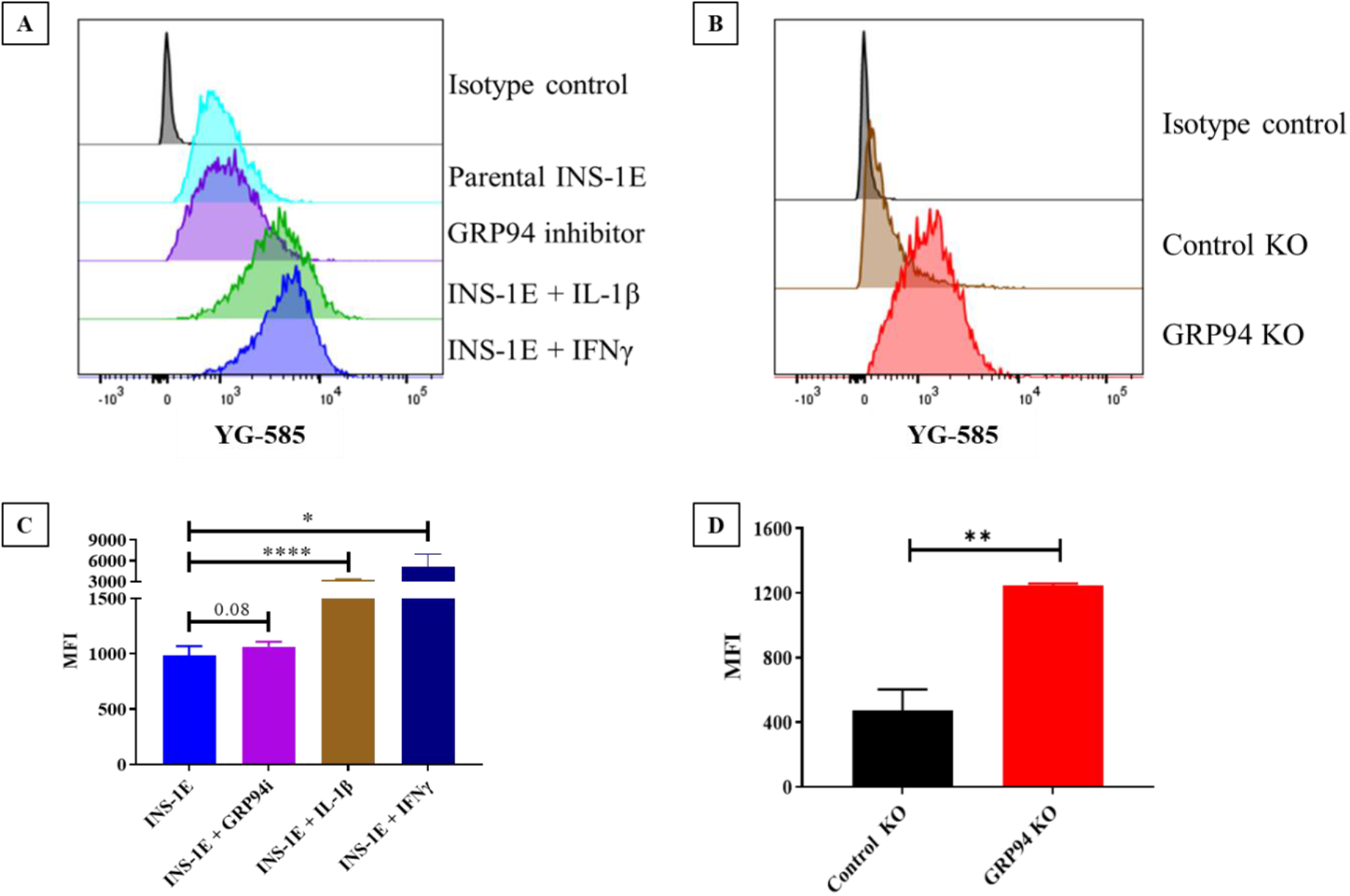
Flow cytometry comparison for RT1.A expression in INS-1E cells under tested conditions. (**A & B**) INS-1E cells compared with relevant controls for RT1.A expression under tested conditions and (**C & D**) Mean fluorescence intensity statistical comparison for RT1.A expression in INS-1E cells in tested conditions versus relevant controls. N = 3 for **C** and = 6 for **D**. Data is presented as mean ± SD. MFI = mean fluorescence intensity. Drugs/cytokines dose and exposure: GRP94i = 20µM, IL-1β = 15pg/ml, IFNγ = 10ng/ml for 24 hours.

### 3.3. GRP94 KO modulates RT1.A-bound peptides yield

The unchallenged INS-1E cells or GRP94i treated/cytokines exposed/GRP94 KO INS-1E cells were lysed for immunoaffinity capture of RT1.A. The RT1.A bound complexes were fractionated and then subjected to LC-MS/MS. The immunopeptidomics was carried out in two biological replicates for each group. Taken all groups together, we found 4505 peptides (from 1653 proteins, listed in supplementary figure 1, supplementary tables 3 A – C) overlapping or unique to different tested conditions.

Some rat strains, for example AVN carrying RT1.A^a^ haplotype, can present peptides of 8-15 mers (Stevens, et al. 1998). As no previous information is available for INS-1E cells derived from NEDH rats carrying RT1.A^g^ haplotype (le Rolle, et al. 2000), we included the same lengths as for AVN rats in our analysis. The majority of peptides (83 – 90 %) in each group ranged between 8 – 15 mers, confirming minimal background interference in the procedure. The number of peptides obtained ranged from 883 in INS-1E (from 484 source proteins) being the lowest to 3365 peptides (from 1434 source proteins) in IFNγ exposed INS-1E (figure 2-A). The GRP94 KO group showed intermediate yield with 1652 peptides (from 862 source proteins) between unchallenged and cytokines exposed INS-1E (figure 2-A). Out of 1652 peptides found in the GRP94 KO groups, 1063 peptides (from 659 source proteins), 75% were overlapping between the two clones confirming low clonal variation. The peptide overlap between replicates for each tested group ranged between 50 – 75 % (figure 2 B).

**Figure 2.**
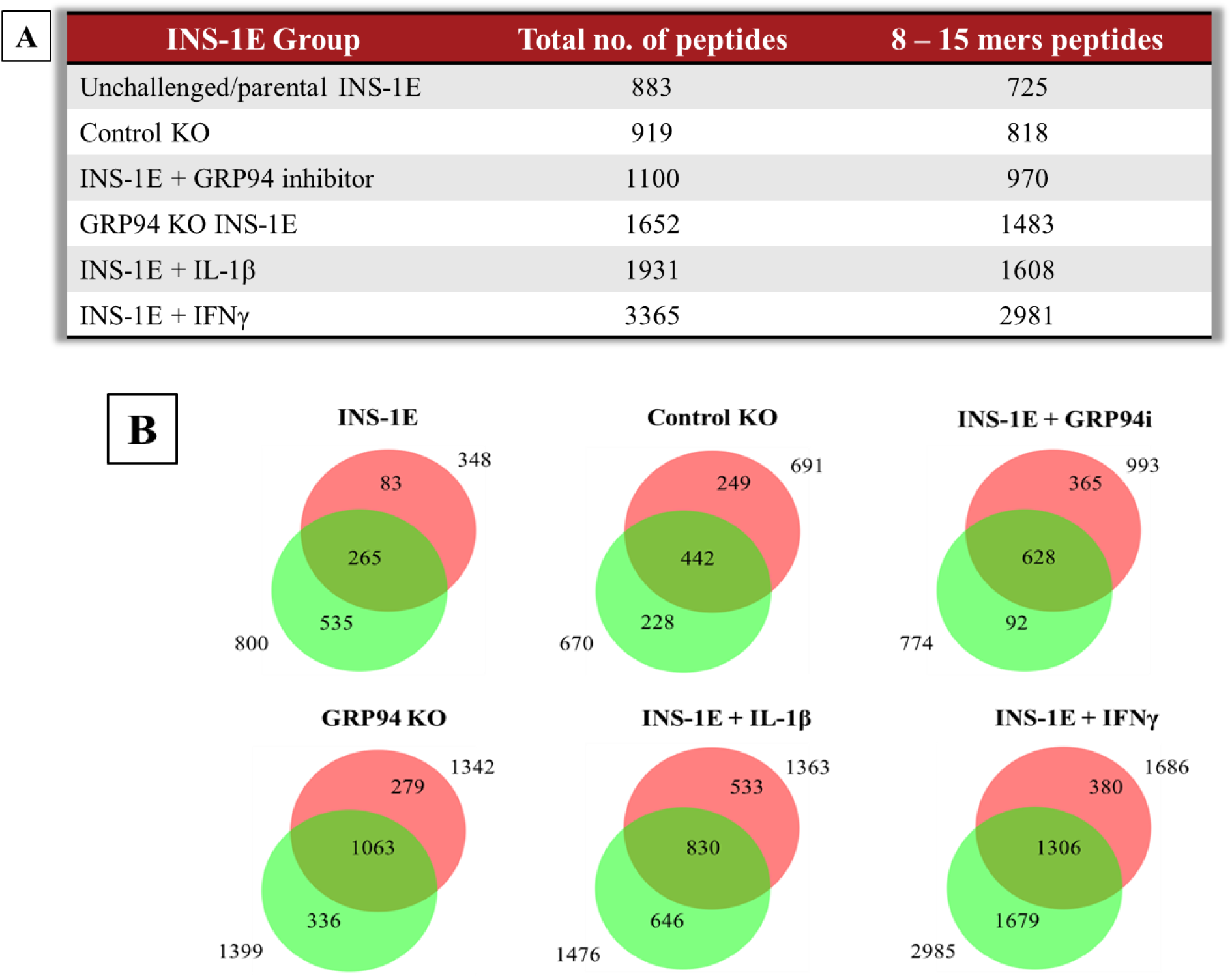
**(A)** Summary table of RT1.A-bound 8 – 15 amino acids length peptides from INS-1E cells under tested conditions. **(B)** Venn diagrams showing peptides overlap between the individual replicates per condition. N = 2 for all groups except for GRP94 KO where N = 4

To exclude peptides present in control conditions, we subtracted peptides present in unchallenged INS-1E and control KO groups from GRP94 KO, IL-1β or IFNγ exposed groups (figure 4 A-C). We found 362 peptides (from 206 proteins) unique to GPR94 KO INS-1E cells compared to unchallenged INS-1E and control KO, while IL-1β or IFNγ exposed INS-1E had 991 and 2416 unique peptides, respectively (figure 3 A-C). We then compared GRP94 KO and cytokines exposed groups to obtain GRP94 KO exclusive and inflammatory setting exclusive peptides. Fifty-three of these peptides were exclusive to GRP94 KO, while 346 and 1726 peptides were unique to IL-1β or IFNγ exposed groups, respectively (figure 3-D). The overlap between GRP94 and cytokines exposed group peptides suggests some similarity between limited folding capacity and inflammatory response. We also found peptides from other proteins localized in the secretory granules between GRP94 KO and cytokines groups e.g. of islet amyloid polypeptide (IAPP) and chromogranin-A (CMGA). Altogether, these data indicates the ER stress and inflammatory stress partly overlap in their activity to modulate MHC-I peptidome.

**Figure 3.**
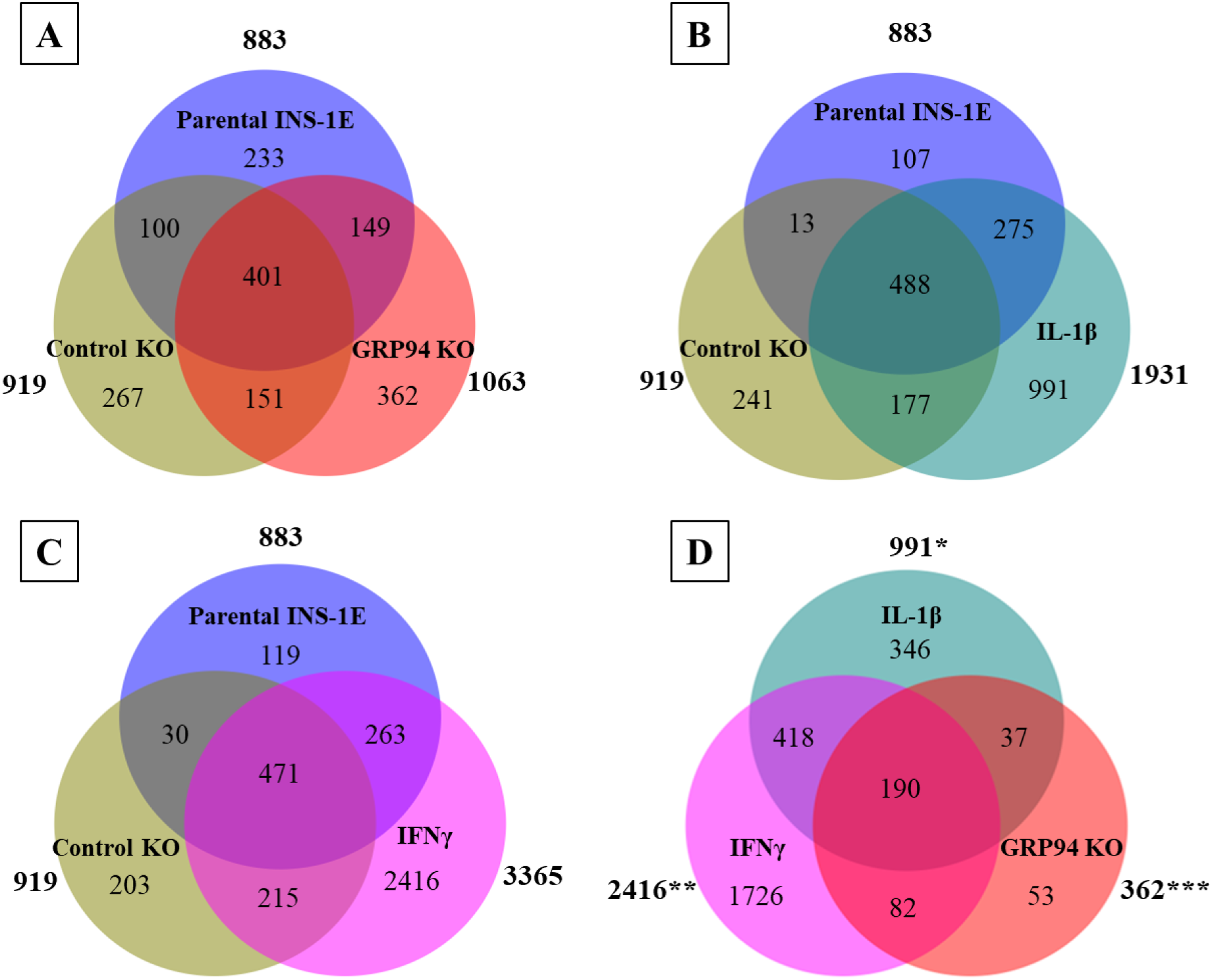
Venn diagrams showing number of MHC I-eluted peptides distinct or overlapping between **(A)** Parental INS-1E, control KO & GRP94 KO and **(B)** Parental INS-1E, control KO & IL-1β exposure (15 pg/ml for 24 hours) **(C)** Parental INS-1E, control KO & IFNγ exposure (10 ng/ml for 24 hours) and **(D)** Peptides exclusive for IL-1β, IFNγ exposed and GRP94 KO INS-1E groups. N = 2 for all except GRP94 KO where N = 4 * = IL-1β peptides excluding parental INS-1E and control KO clone ** = IFNγ peptides excluding parental INS-1E and control KO clone *** = GRP94 KO peptides excluding parental INS-1E and control KO clone

**Figure 4.**
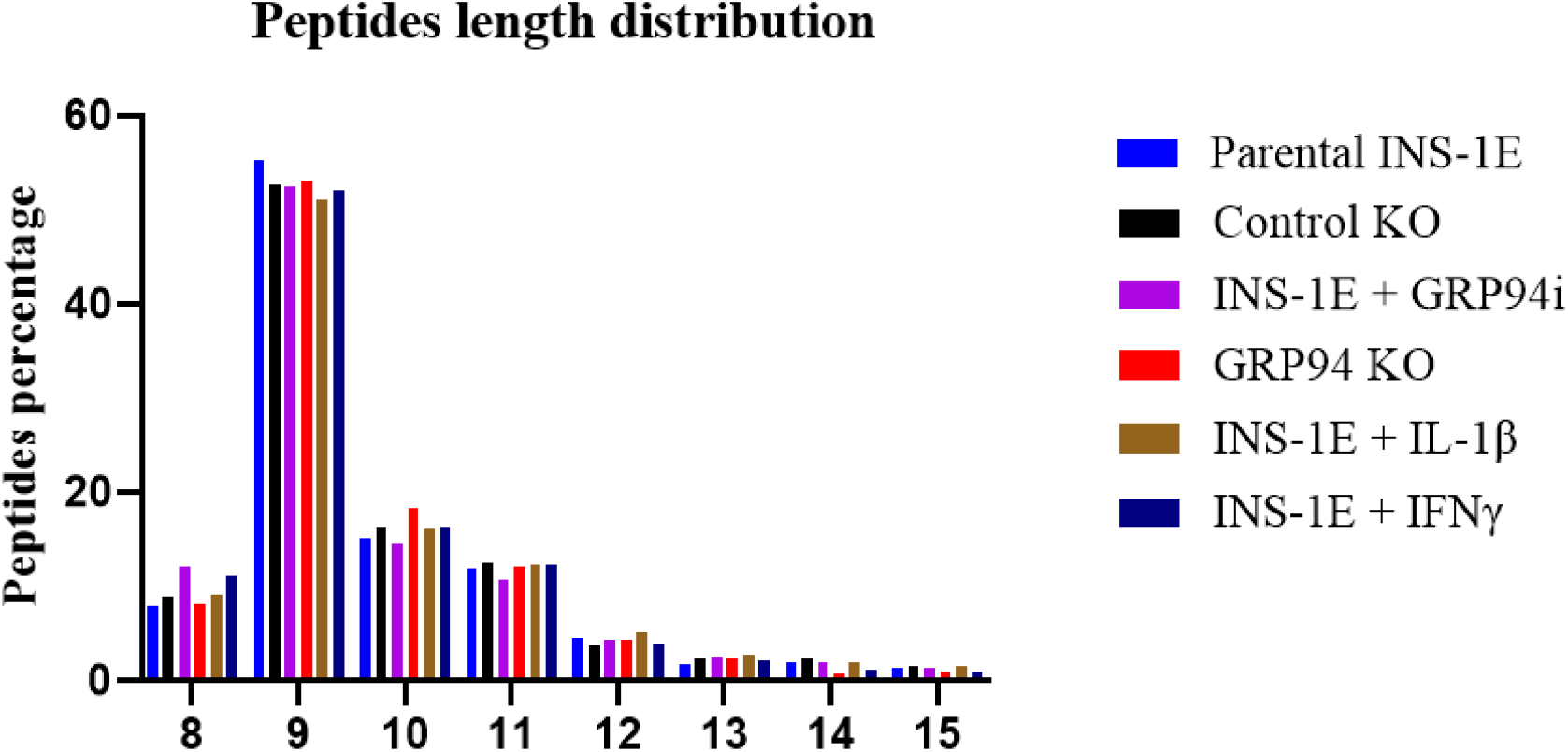
Length distribution for 8 – 15 amino acid-length peptides between parental INS-1E, control and confirmed GRP94 KO INS-1E, 24 hour treatment with GRP94 inhibitor (20 µM), or INS-1E cells exposed to cytokines IL-1β (15 pg/ml) or IFNγ (10 ng/ml). N = 2 for all except GRP94 KO where N = 4.

### 3.4. RT1.A^g^ has high affinity for nonamers

The length distribution showed nonamers to comprise the highest proportion (around 55 %) in all groups, followed by decamers and undecamers as the second and third most preferred length of peptides for RT1.A^g^ binding, accounting for ∼15 % to 18 % and ∼11 % to 12.5 %, respectively in the different groups (figure 4), suggesting a minimal variation among all tested conditions.

The peptide-binding motifs showed high affinity for leucine, isoleucine and valine at P2 while arginine and lysine dominated PΩ, (figure 5 & 6). No obvious differences were observed in amino acid preferences within the peptides in any group. The only exception was seen in IFNγ exposed INS-1E cells where tyrosine appeared as the third most abundant amino acid at PΩ in nonamers (figure 5 & 6).

**Figure 5.**
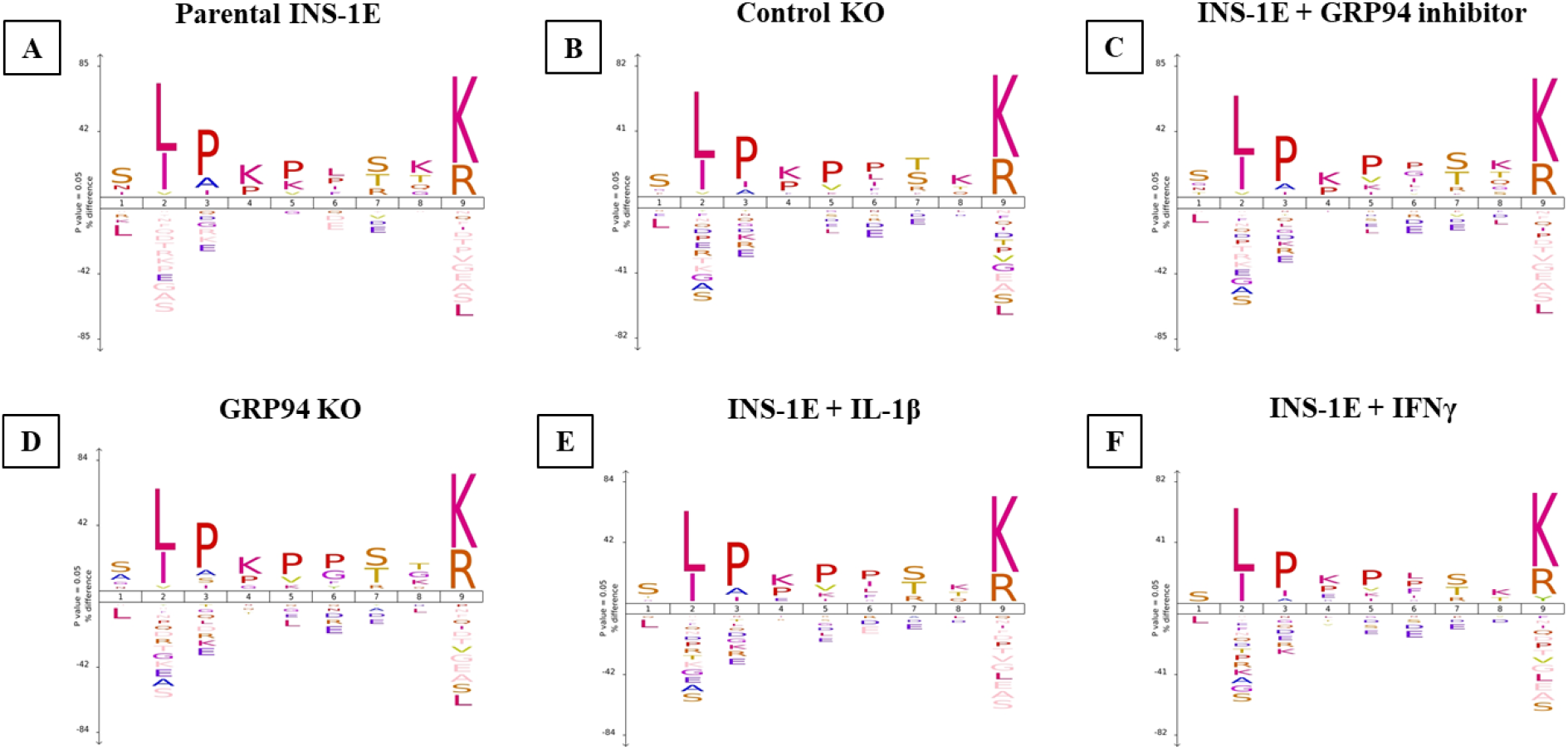
Nonamers binding motif for INS-1E cells under tested conditions. **(A)** parental INS-1E **(B)** control GRP94 KO clone **(C)** INS-1E treated with GRP94 inhibitor (20 µM for 24 hours) **(D)** GRP94 KO INS-1E clone **(E)** IL-1β exposure (15 pg/ml for 24 hours) and **(F)** IFNγ exposure (10 ng/ml for 24 hours). N = 2 for all except GRP94 KO where N = 4.

**Figure 6.**
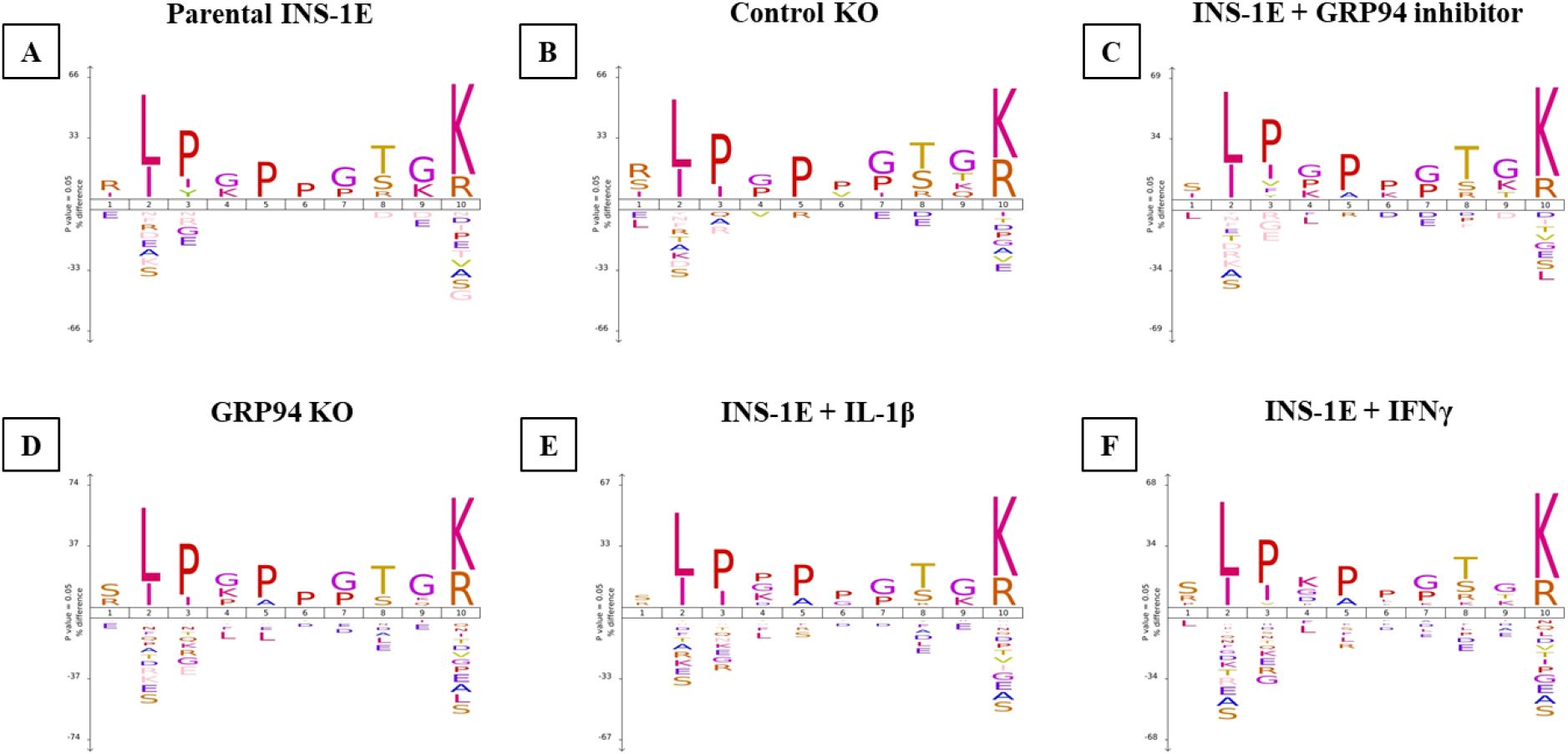
Decamers binding motif for INS-1E under tested conditions. **(A)** parental INS-1E **(B)** control GRP94 KO clone **(C)** INS-1E treated with GRP94 inhibitor (20 µM for 24 hours) **(D)** GRP94 KO INS-1E clone **(E)** IL-1β exposure (15 pg/ml for 24 hours) and **(F)** IFNγ exposure (10 ng/ml for 24 hours). N = 2 for all except GRP94 KO where N = 4.

The Gibbscluster analysis for peptides from all tested groups also confirmed high turnover of peptides with similar binding motif (supplementary figure 2).

### 3.5. GRP94 KO/cytokines exposure modifies proinsulin derived RT1.A-bound peptides

In addition to the general increase in peptide repertoire in GRP94 KO and cytokines exposed INS-1E cells, we also found a qualitative increase in proinsulin derived peptides. Three peptides derived from proinsulin were found in one or both cytokines group (VLWEPKPAQAFVK, ILWEPKPAQAFVK, ALWMRFLPL) (table 1). Moreover, we also found four peptides (HLVEALYL, ALYLVCGERGFF, ALYLVCGERGFFYTP, and YLVCGERGFF) common to GRP94 KO/i and cytokines exposed groups. The overlap between cytokines and GRP94 KO/i groups suggest a possible interplay between defective folding and inflammatory pathways. Interestingly, we found two (pro) insulin peptides FFYTPKS and VEDPQVPQ unique to GRP94 KO/i groups. The peptide FFYTPKS is a non-canonical 7-mer but was detected in both GRP94 KO/i conditions.

**Table 1.**
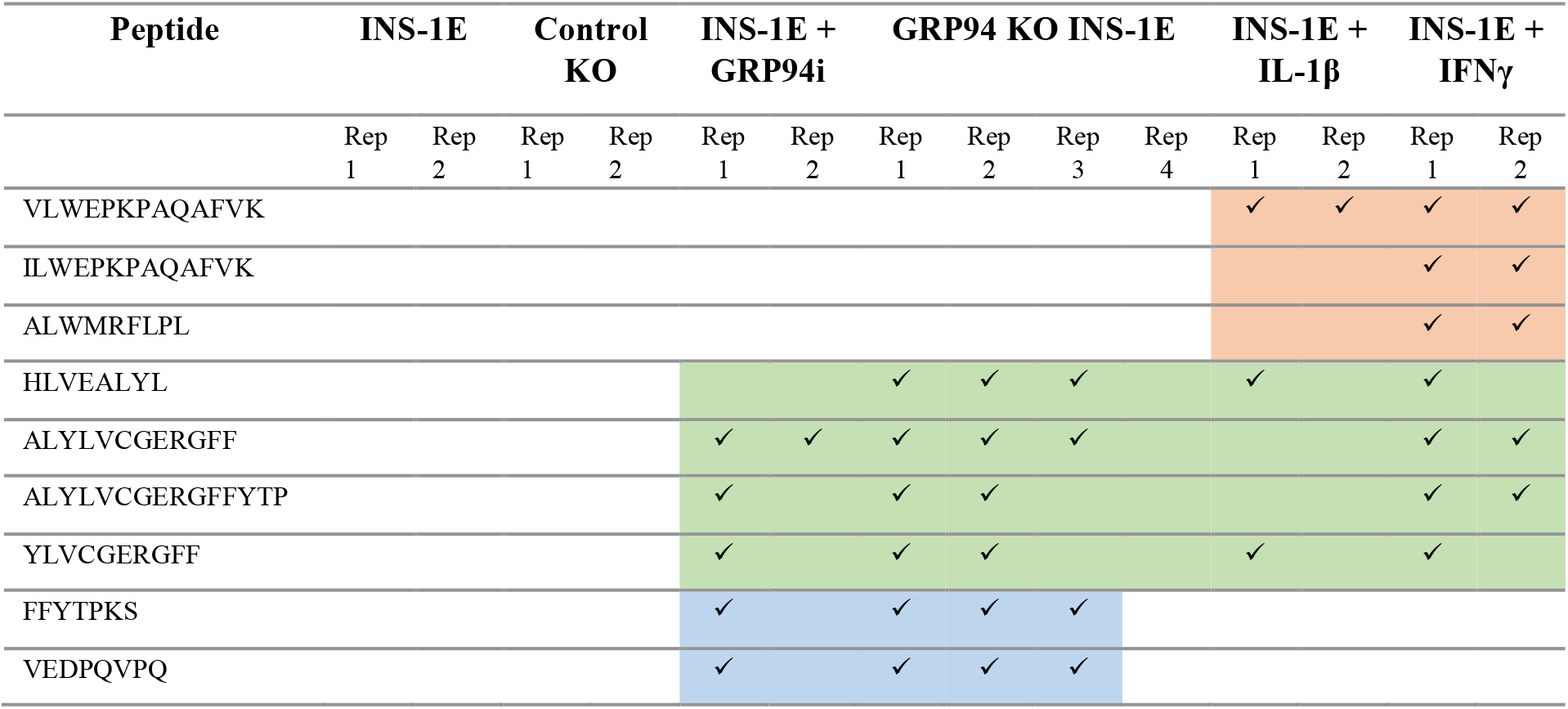
RT1.A eluted proinsulin peptides from INS-1E cells derived from (pre)-proinsulin overlapping or unique for GRP94 KO/KD or cytokines exposure. N = 2 for all groups except GRP94 KO where N = 4.

We trained the artificial neural networking platform NNAlign to scan insulin sequences for occurrence of binding motifs and to order them as potential high scoring binders among INS-1E-derived peptides from all groups excluding the proinsulin partite. The binding motif came out similar to Gibbscluster analysis (supplementary figure 3). With the exception of GRP94i group, we found increased peptide yield in treatment groups (Supplementary table 4) with higher scores compared to control but lower than best peptides from IFNγ exposed group. The peptide ALYLVCGERGFF, corresponding to insulin chain B:14–25 appeared as high scoring peptide in GRP94 KO/i and IFNγ exposed group. HLVEALYL (B:10–B17) appeared as top scoring peptide GRP94 KO and IL-1β group as well as second highest scoring peptide in IFNγ group. GFFYPTMS (B:23–30) was the highest scoring peptide in IFNγ group with a score of 0.741. Collectively, these observations indicate an increased likelihood of proinsulin peptides from treatment groups as RT1 binders.

### 3.6. Limiting β-cell folding capacity activates inflammatory pathways

To test whether a defect in proinsulin folding capacity in β-cells leads to the activation of inflammatory pathways, we tested expression of the NALP1, IκBα and IL-1β content and secretion in our β-cell model. We found that NALP1 expression (at 165 kDa fully organized NALP1 and 70 kDa corresponding to NALP1 lacking leucine rich repeats (LRR)) was upregulated in GRP94 KO cells compared to control group (figure 7-A-C). Typically NALP1 consists of leucine rich repeats (LRR), a NACHT domain and a PYD/CARD domain. In general LRR is considered to play role as ligand recognition domain for inflammasomes; however, a recent study showed NLRP3 are active despite lacking LRR and thus LRR is dispensable (Hafner-Bratkovič, et al. 2018). The IκBα on the other hand was found to be decreased in GRP94 KO group (figure 7-A&D). We also determined cell contents of proIL-1β, a canonical substrate of NALP1 processing. In accordance with the increased NALP1 expression, we found that pro-IL-1β content was reduced in GRP94 KO compared to control KO while mature IL-1β content was hardly detectable but increased in secretion (figure 7-A&E-F). Taken together, these data suggest that limited β-cell folding capacity is associated with NFκB activation, NALP1 expression and activity, proIL-1β processing and secretion of IL-1β.

**Figure 7.**
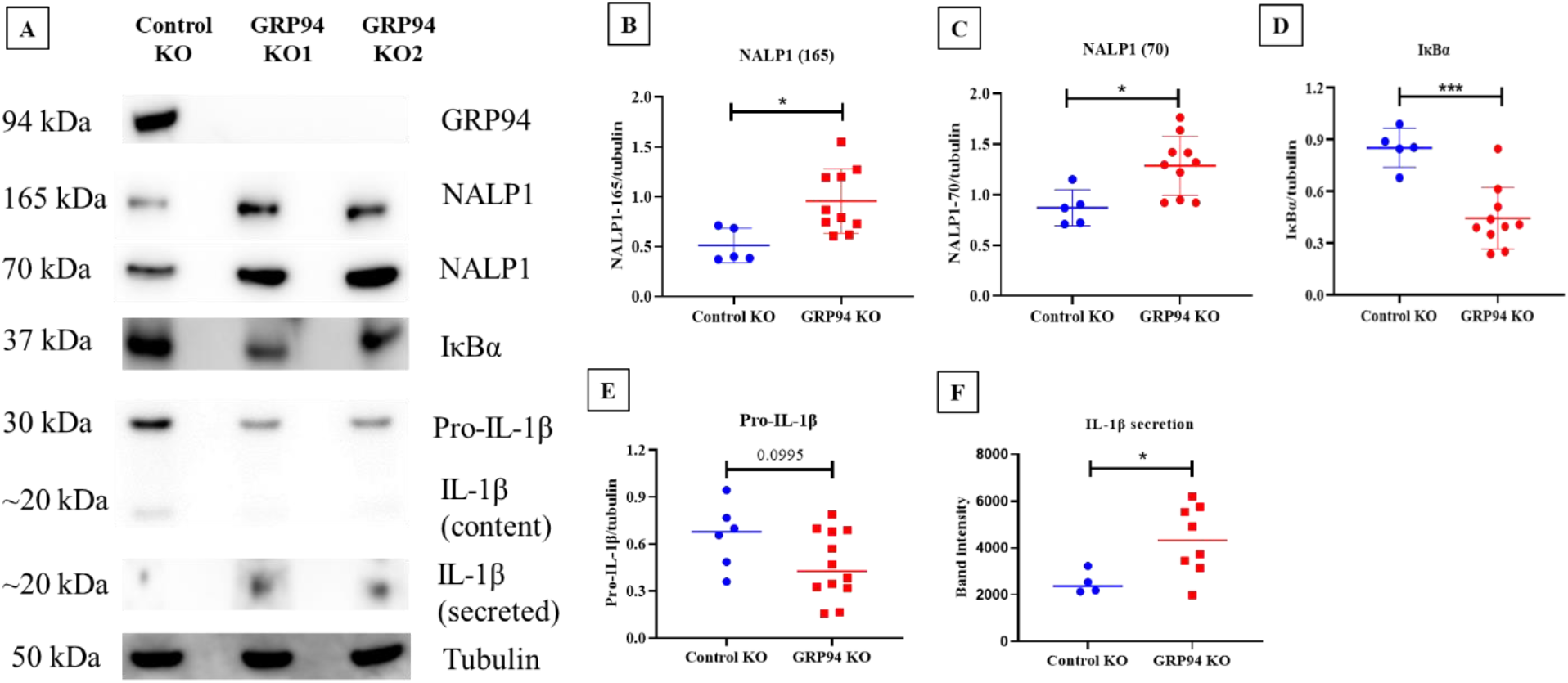
Western blot analysis for the expression of NALP1, IκBα and IL-1β in GRP94 KO INS-1E and control KO cells. **(A)** SDS-PAGE for expression of NALP1 (fully assembled version (165 kDa) and without LRR (70 kDa)), IκBα and IL-1β in tested conditions. (**B-F)** Statistical analysis (unpaired t-test) for the expression of NALP1, IκBα and pro-IL-1β/IL-1β quantified and normalized to tubulin in tested conditions. N = 4 – 6 for different proteins of interest.

## 4. Discussion

The long-standing knowledge gap in understanding T1D pathophysiology is how tolerance is lost against β-cell self-peptides, resulting in immune mediated destruction of β-cells. In this study, we provide the proof-of-principle that ER stress as a result of diminished or functional-loss of β-cell folding capacity leads to an increased expression of the MHC-I self-antigen presentation platform, a modulation of the MHC-I bound peptidome as well as activation of inflammatory pathways. We also established the first database for haplotype RT1.A^g^ bound peptides endogenously processed and presented in one of the most commonly used β-cell line, INS-1E, derived from NEDH rats.

The ER has a key role in MHC class I antigen presentation pathway as peptides generated by proteasome are transported into the ER and loaded onto MHC class I for immune surveillance (Blum, et al. 2013). Protein misfolding in the ER, e.g. tyrosinase misfolding has been shown to increase MHC antigen presentation efficiency (Ostankovitch, et al. 2005). MHC I hyper expression is a hallmark of T1D as seen in islets from newly diagnosed patients (Coppieters, et al. 2012; Richardson, et al. 2016) and an elevated MHC I expression facilitates antigen presentation to CD8^+^ T-cells (Pugliese 2017). We found an increased RT1 expression in GRP94 KO cells as well as upon cytokines exposure (figure 1-A – D). The increased RT1 expression may be mediated via NF-κB pathway, which itself is activated by ER stress response or cytokines (Ghemrawi, et al. 2018; Salvado, et al. 2015). Indeed, GRP94 KO INS-1E cells exhibit PERK/eIF2α activation (Ghiasi, et al. 2019) which may attenuate IκBα translation (Deng, et al. 2004) (figure 7). This could lead to the activation of NF-κB to promote transcription of proinflammatory genes and upregulate MHC-I expression (Forloni, et al. 2010). The increased MHC-I expression in the islet microenvironment due to ongoing inflammatory processes (Marroqui, et al. 2017) makes β-cells more visible to the effector T-cells and possibly explains β-cell vulnerability in T1D patients despite the similar frequency of circulating β-cell reactive CD8^+^ T-cells in both T1D patients and healthy individuals (Culina, et al. 2018; Gonzalez-Duque, et al. 2018).

We found that GRP94 KO/i increased RT1-bound peptide repertoire as did the cytokines exposure (figure 2-A & B). The GRP94 KO group peptide repertoire overlapped partly with those from cytokine exposed groups (figure 3-A – D) suggesting some shared signaling pathways between these two separate inducers of ER stress. Interestingly, the GRP94 KO also shared (pro) insulin peptides with the cytokines group. (Pro) insulin peptides especially those derived from proinsulin B-chain are recognized as neoepitopes and listed e.g. in (James, et al. 2020). We have found four peptides (HLVEALYL, ALYLVERGERGFF, ALYLVCGERGFFYTP and YLVCGERGFF) from insulin B-chain (table 1) common in GRP94 KO and cytokines exposed experimental groups. Diminished GRP94 activity is expected to affect proinsulin processing (Ghiasi, et al. 2019) as do the cytokines such as IL-1β (Spinas, et al. 1986), suggesting ER stress induced by limited folding capacity mimics the action of cytokines on (pro) insulin processing. The two GRP94 KO/i exclusive insulin peptides are VEDPQVPQ (C-peptide: 2 – 9) and FFYTPKS (B: 25 – 31) (table 1). As defect in GPR94 activity leads to mishandling at proinsulin level, it is plausible that C-peptide also contributed to MHC-I peptidome in GRP94 KO group. The FFYTPKS is a non-canonical peptide but appeared in majority of mass spec analyses, however despite proper control set up it may still be a non-specific binder. The presence of other β-cell proteins such as IAPP (ILVALGHLR) and CMGA (RMQLAKELT) overlapping in both GPR94 KO and cytokine exposed groups (supplementary table 3.a) suggests that the GRP94 functional deficiency impact on protein folding in ER might not be restricted to proinsulin and warrants further investigations.

Studies show ER stress precedes inflammation as well as the development of T1D (Hasnain, et al. 2012) (Tersey, et al. 2012). The interesting question is how in our experimental settings, limited folding capacity coupled with compensated UPR activation (Ghiasi, et al. 2019) activates inflammation pathways? As our experiments were conducted in an *in vitro* settings, the cells must be reliant on internal mechanisms to activate inflammatory pathways. As PERK has been reported to increase inflammasome expression (D’Osualdo, et al. 2015), we tested expression levels of NALP1 and its downstream targets, IL-1β and IκBα in our β-cell model. Interestingly, we found an increased expression of NALP1 in GRP94 KO group compared to control group (figure 7). Typically, the inflammasome activation requires two signals. Signal-1 such as glucose, lipopolysaccahrides or cytokines (Bauernfeind, et al. 2009; Schroder, et al. 2012; Wang, et al. 2020) is the priming signal that upregulates inflammasome and increases proIL-1β expression in response to ligand binding the toll-like receptors (TLR), whereas signal-2 provided by extracellular ATP, toxins and particulate matters such as IAPP (Kelley, et al. 2019; Mariathasan, et al. 2006; Masters, et al. 2010) activates the inflammasome-dependent activation of caspase-1, which in turn cleaves proIL-1β and proIL-18 into mature and biologically active IL-1β and IL-18. Some drugs that induce ER stress such as thapsigargin and tunicamycin or mitochondrial oxidative stress inducers like rotenone can provide both signal-1 and signal-2 via promoting interaction of NALP with thioredoxin-interacting protein (TXNIP) (Lerner, et al. 2012; Wali, et al. 2014; Zhou, et al. 2010; Zhou, et al. 2011). PERK activation in GRP94 KO cells can lead to increased inflammasome activation (D’Osualdo, et al. 2015) via thioredoxin interacting protein (TXNIP) as in (Lerner, et al. 2012). To confirm NALP1 overexpression is followed by inflammasome activation, we estimated levels of (pro) IL-1β content as well as secretion. We found decreased pro-IL-1β content in GRP94 KO cells possibly due to secretion as mature IL-1β. Indeed we found a high amount of IL-1β in the concentrated supernatants of INS-1E cells confirming high activity of NALP1 inflammasome in GRP94 deficient β-cells and possibly facilitate the onset of inflammatory events in the islet microenvironment.

The importance of GRP94 in β-cells has gained the attention only recently. GRP94 has long been known as critical for the prenatal development and ER quality control (Marzec, et al. 2012) as is the case with pancreas (Kim, et al. 2018). Conditional knockout of GRP94 in mice leads to pancreatic hypoplasia and reduced β-cell mass and proliferation (Kim, et al. 2018). Islets derived from type 2 diabetes patients showed reduced GRP94 levels (Ghiasi, et al. 2018). Moreover, plasma of T1D patients contains elevated concentrations of free GPR94 as well as circulating GRP94-bound complexes such as GPR94-α1–antitrypsin (Pagetta, et al. 2003) and GRP94-IgG (Pagetta, et al. 2007; Roveri, et al. 2015). The extracellular GRP94 represents immunological danger as it stabilizes the bound peptide (Berwin, et al. 2001; Calderwood, et al. 2007) and increases the efficiency of uptake by scavenger receptor type A (SR-A) and CD91 for cross-presentation (Basu, et al. 2001; Berwin, et al. 2002a; Berwin, et al. 2002b; Binder, et al. 2000). It may therefore be of interest to explore the presence of GRP94 bound to its clientele, such as GRP94-PI complex, in circulation in T1D patients as potential therapeutic marker.

Another study showed an increased macrophage polarization, accompanied by inflammatory environment and development of insulin resistance in conditional GRP94 KO in macrophages in mice (Song, et al. 2020), suggesting a connection between GRP94 and associated molecules in facilitating inflammatory response and should be tested also in β-cells with loss of function for GRP94. Another element that bridges ER stress to inflammation are viral infections as cells are burdened with massive synthesis of viral proteins, which may result in an increase of endogenous proteins misfolding (He 2006) as well as induction of proinflammatory environment (Dotta, et al. 2007).

A third possible connection may lie between UPR and inflammation. Activation of classical UPR is known to promote inflammatory signaling (Hasnain, et al. 2012). PERK/eIF2α activation mediated IκBα translational inhibition (Deng et al. 2004) or rapid proteasomal degradation (Mathes, et al. 2008) may decrease IκB/ NF-κB ratio and facilitate nuclear translocation of NF-κB as discussed in the previous paragraph. So theoretically, GRP94 KO induced ER stress and inflammation can intersect at multiple levels and my lead to inflammatory settings that trigger the immune response against β-cells as proposed in figure 8.

**Figure 8.**
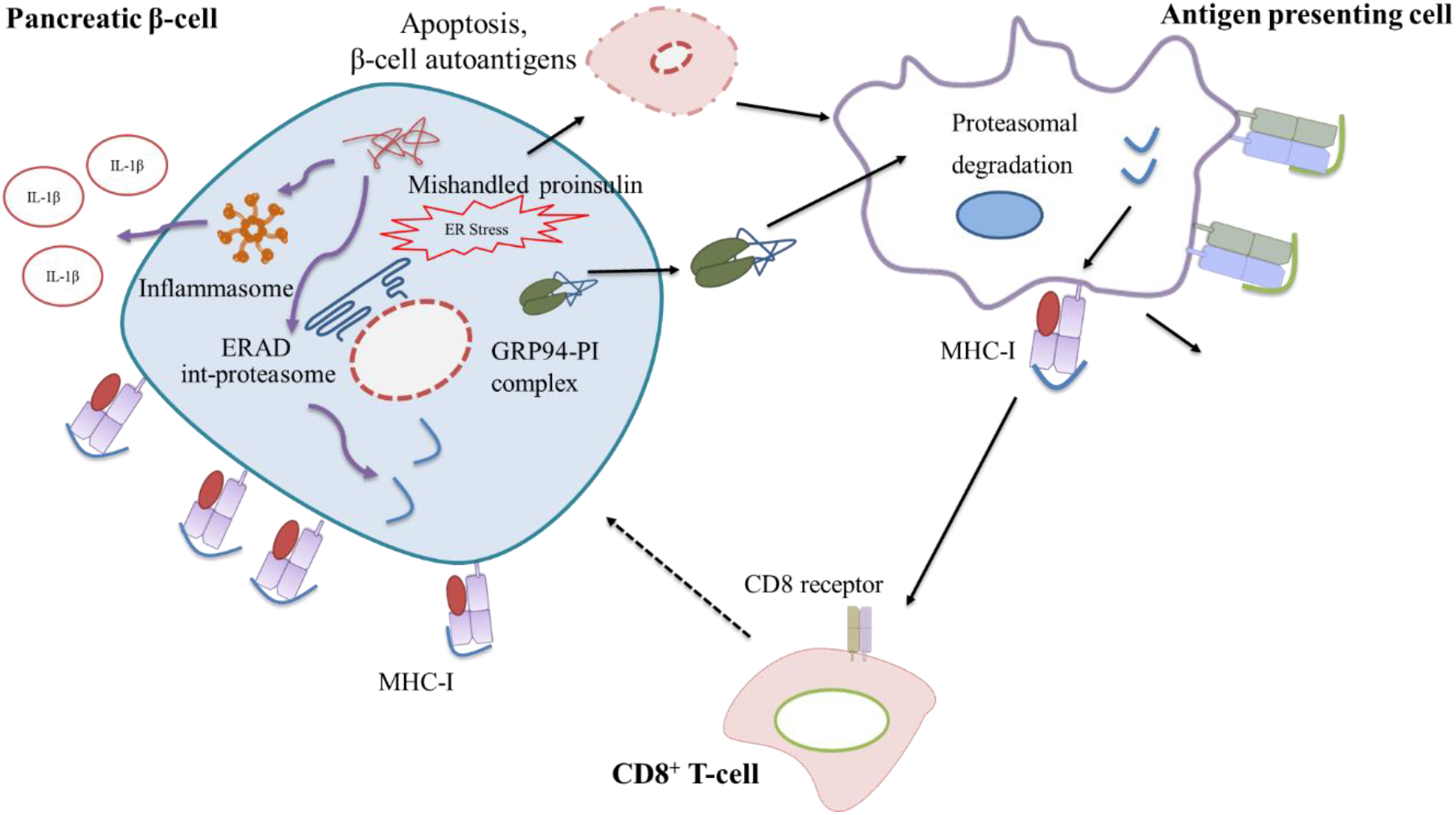
Proposed mechanism of autoimmunity induction in T1D. A defect in the ability of GRP94 to process proinsulin leads to an accumulation of mishandled proinsulin in β-cells. This leads to ER stress, activation of inflammatory pathways, assembly of int-proteasome and secretion of GRP94-PI complexes. Simultaneously the increased MHC-I expression on β-cells make them more visible to immune cells. The secreted GRP94-PI complexes are taken up by APCs and cross presented to CD8^+^ T-cells to trigger immune attack against β-cells.

## 5. Limitations of the study/Further possibilities

The role of GRP94 KO/cytokines specific (pro) insulin peptides we found in our immunopeptidomics analysis in the induction of autoimmunity needs to be investigated in patients as neoepitopes. A preliminary further analysis could have been made using bioinformatics tools such as NetMHC (Andreatta and Nielsen 2016) that uses trained artificial neural networks to make predictions for strong MHC class-I binding. However, this tool is not trained/optimized for rat strains-based haplotypes due to literature scarcity. This is especially true for INS-1E cells, which carry RT1.A^g^ haplotype and no previous report is available for comparison. In addition, the IEDB database lists only one MHC class-II epitope (FVKQHLCGSHLVEALYLV) derived from insulin in BB rats having RT1.A^u^ haplotype (Heath, et al. 1999). In addition, this approach of ER stress induced by limited β-cell folding capacity leading to activation of inflammatory pathways should be expanded to e.g. rodent models of diabetes such as NOD mouse/BB rat with tissue specific GRP94 KO/targeted mutation or ER stress induced by thapsigargin in human cell lines/islets.

One more constraint in expanding our analysis is related to less extensive proteome explored for rats (8,133 reviewed proteins) compared to mouse (17,085 reviewed entries) or human (20,360 reviewed entries) on Uniprot until 30^th^ November 2021 to compare. Therefore, it is reasonable to deduce that we actually detected more RT1-bound peptides in our experimental settings but could not identify them in the database due to a relatively limited rat proteome data availability.

## 6. Conclusion

In conclusion, we found that β-cells challenged with a defect in proinsulin folding capacity lay the groundwork for increased MHC I presentation of altered peptidome. Furthermore, this happens via activation of inflammatory pathways in β-cells, sensitizing them for immune attack. Further experimental validation of our proposed idea in more models are needed to advance this concept for translational benefits.

## Supplementary materials

Supplementary materials with list of peptides/source proteins and NNAlign table can be accessed via request to.

## Author contributions

Conceptualization: M.T.M., T.M.-P., M.S.K., A.W.P., P.F., J.K. ; methodology: M.S.K., T.D. E.P.-M., R.A.; formal analysis: M.S.K., M.T.M., T.M.-P., A.W.P., P.F., J.K., S. B., M.N. ; Resources: M.T.M., T.M.-P., A.W.P. ; data curation: M.S.K., T.D., E.P.-M., R.A. ; manuscript preparation: M.S.K., M.T.M., T.M.-P., A.W.P., P.F., R.A., E.P.-M., J.K., S. B., M.N. ; supervision: M.T.M., T.M.-P., A.W.P., P.F.

## Acknowledgement

The authors would like to thank Professor Claes Wollheim (University Medical Center, Geneva, Switzerland) for generously sharing INS-1E cells.

---

**Supplementary table 1.** List of chemicals and antibodies used for experimental work.

| **Reagent** | **Commercial vendor, Cat#** |
| --- | --- |
| **Ethanol** | Ajax Finechem, cat. no. AJA214 |
| **Optima water** | Thermo Fisher Scientific, cat. no. FSBW6-4 |
| **Methanol** | Merck Millipore, cat. no. 1.06018.4000 |
| **Acetonitrile** | Thermo Fisher Scientific, cat. no. FSBA955-4 |
| **Formic acid** | Sigma-Aldrich, cat. no. 14265-1ML |
| **Acetic acid** | Sigma-Aldrich, cat. no. 33209-1L-GL |
| **Igepal CA-630** | Sigma-Aldrich, cat. no. I8896 |
| **Tris** | Astral Scientific, cat. no. BIO3094T |
| **Antibodies** |  |
| **Anti-rabbit NALP1** | Cell signaling, cat. no. 4990S, dilution = 1:2,000 |
| **Anti-rat GRP94** | Thermofisher Scientific, cat. no. MA3-016, dilution = 1:10,000 |
| **Anti-mouse proinsulin** | Cell Signaling, cat. no. 8138S, dilution = 1:5000 |
| **Anti-mouse tubulin** | Sigma, cat. no. T6074, dilution = 1:10,000 |
| **Anti-mouse IκB-α** | Thermo Fisher scientific, cat. no. 40903, dilution = 1:1,000 |
| **Recombinant Anti-IL-1β** | Abcam, cat. no. ab283818, dilution = 1:2,000 |
| **Anti-rat** | Abcam, cat. no. ab6734, dilution = 1;10,000 |
| **Anti-rabbit** | Cell signaling, cat. no. 7074S, dilution = 1;10,000 |
| **Anti-mouse** | Cell signaling, cat. no. 7076S, dilution = 1;10,000 |

**Supplementary table 2.**
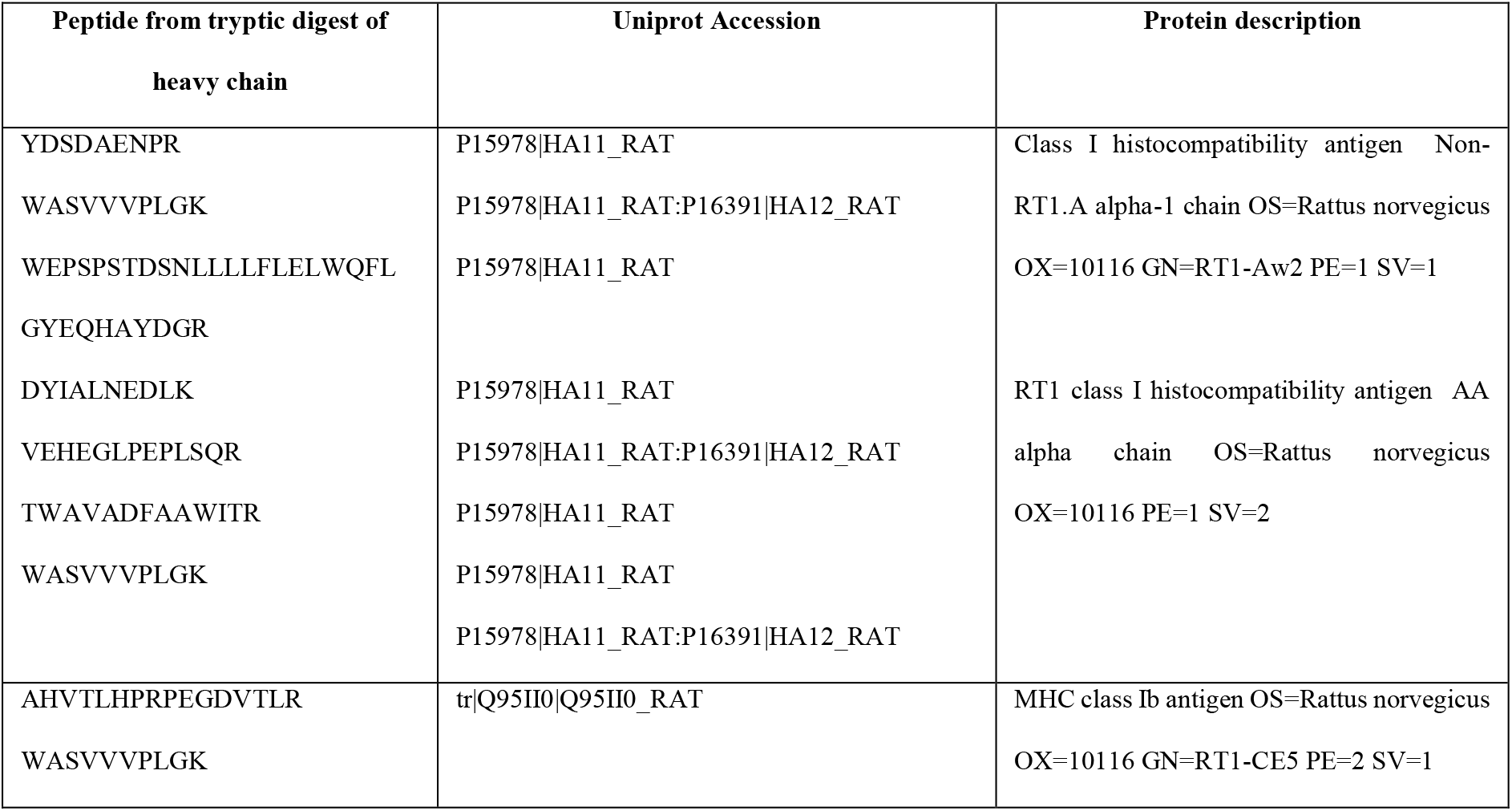
List of peptides from heavy chain fraction digestion of INS-1E cells. The peptides map to both RT1.A (class Ia) and class Ib region (RT1.C/E/M) of RT1 system in *Rattus norvegicus*.

**Supplementary table 3a**. List of RT1.A-bound peptides from individual replicates of INS-1E cells under tested conditions. The data can be accessed via sending email to.

**Supplementary table 3b**. List of peptides combined per condition for all groups. The data can be accessed via sending email to.

**Supplementary table 3c**. List of source proteins for RT1.A bound peptides for all tested groups. The data can be accessed via sending email to.

**Supplementary table 4**. List of peptides with NNAlign predicted score and peptide core for INS-1E cells under tested conditions. The data can be accessed via sending email to.

**Supplementary table 5**. List of all peptides from tryptic digestion of heavy chain fragment. The data can be accessed via sending email to.

**Supplementary Figure 1.**
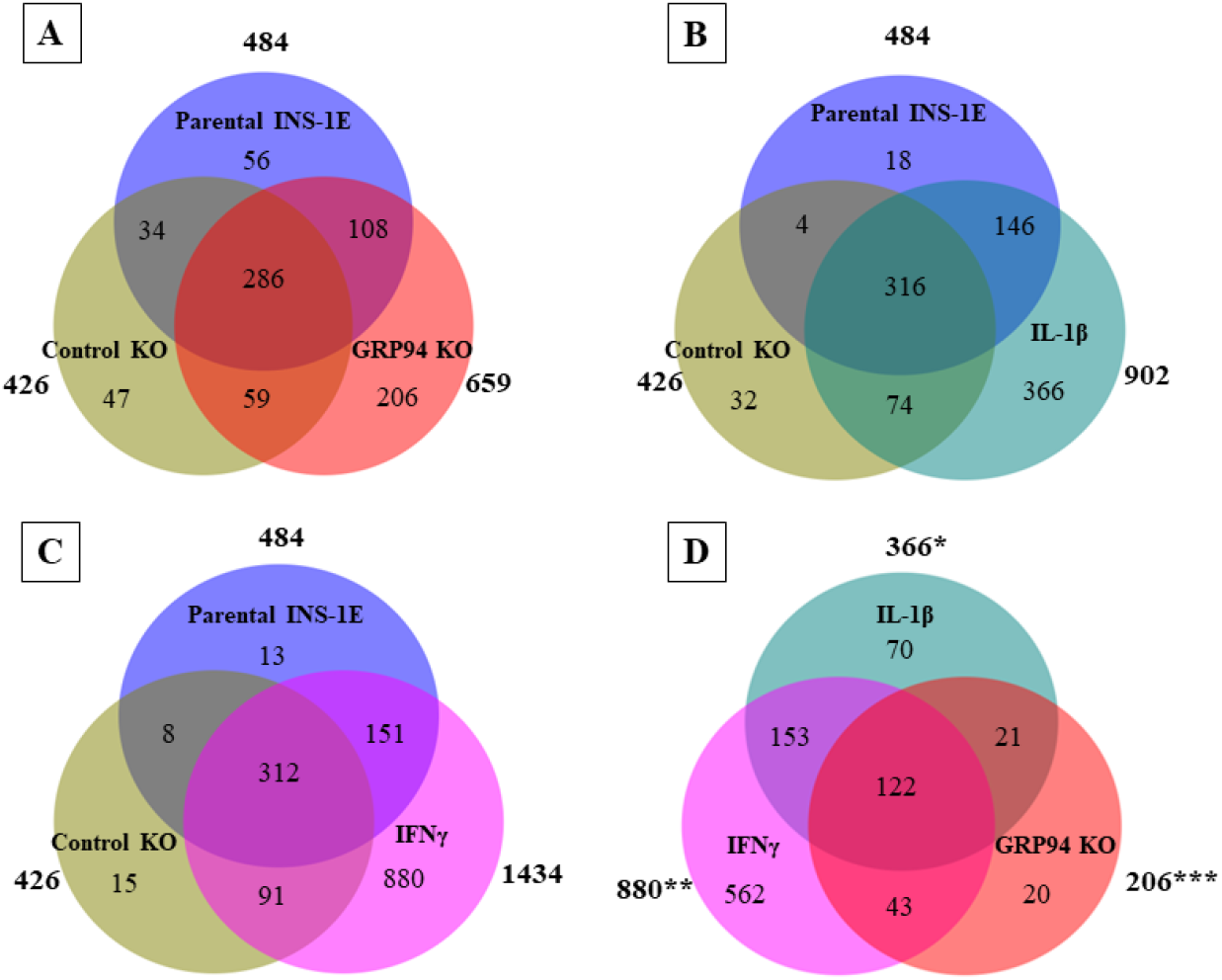
Venn diagrams showing number of source proteins for RT1.A eluted peptides distinct or overlapping between **(A)** Parental INS-1E, control KO & GRP94 KO and **(B)** Parental INS-1E, control KO & IL-1β exposure (15 pg/ml for 24 hours) **(C)** Parental INS-1E, control KO & IFNγ exposure (10 ng/ml for 24 hours) and **(D)** RT1.A contributing proteins exclusive for IL-1β, IFNγ exposed and GRP94 KO INS-1E groups. N = 2 for all except GRP94 KO where N = 4. * = IL-1β group proteins excluding parental INS-1E and control KO clone ** = IFNγ group proteins excluding parental INS-1E and control KO clone *** = GRP94 KO group proteins excluding parental INS-1E and control KO clone

**Supplementary Figure 2.**
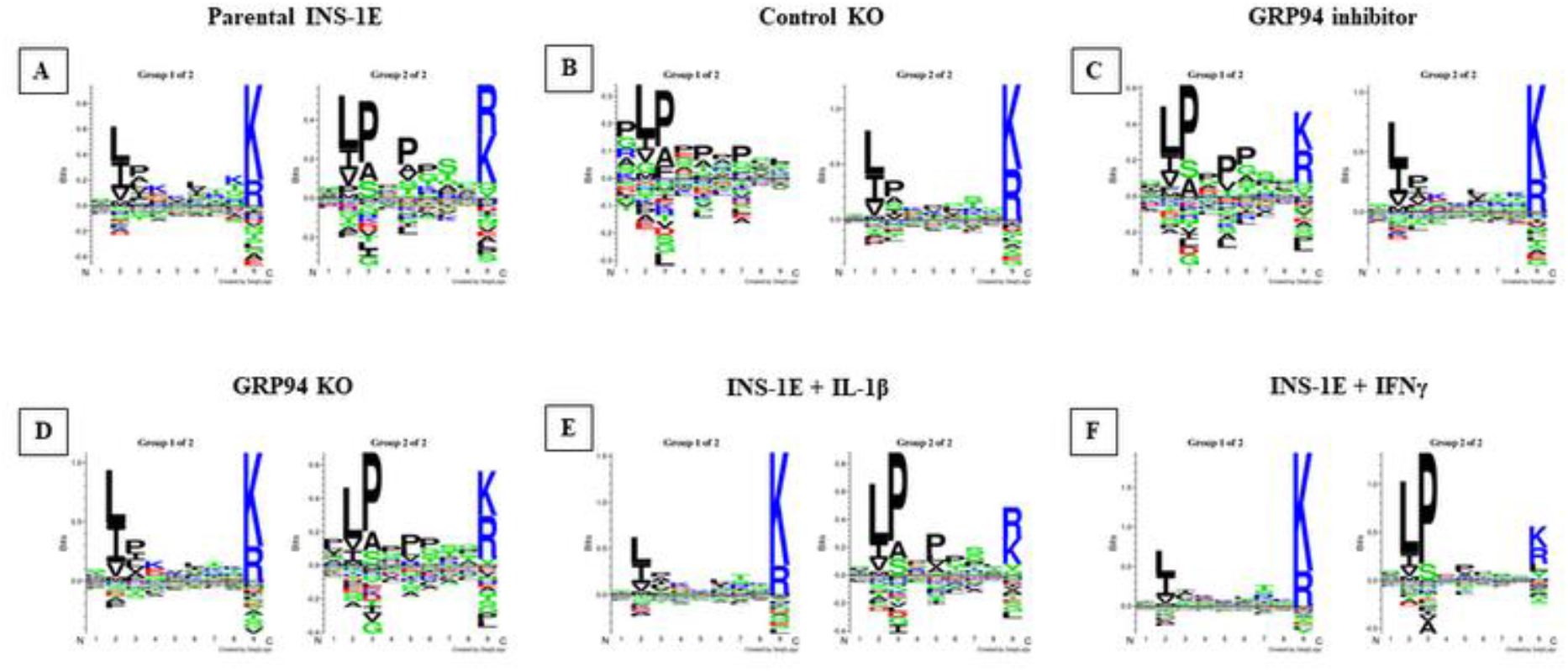
Gibbscluster analysis for RT1.A-bound peptides in INS-1E cells under tested conditions. **(A)** parental INS-1E **(B)** control GRP94 KO clone **(C)** INS-1E treated with GRP94 inhibitor (20 µM for 24 hours) **(D)** GRP94 KO INS-1E clone **(E)** IL-1β exposure (15 pg/ml for 24 hours) and **(F)** IFNγ exposure (10 ng/ml for 24 hours). N = 2 for all except GRP94 KO where N = 4.

**Supplementary Figure 3.**
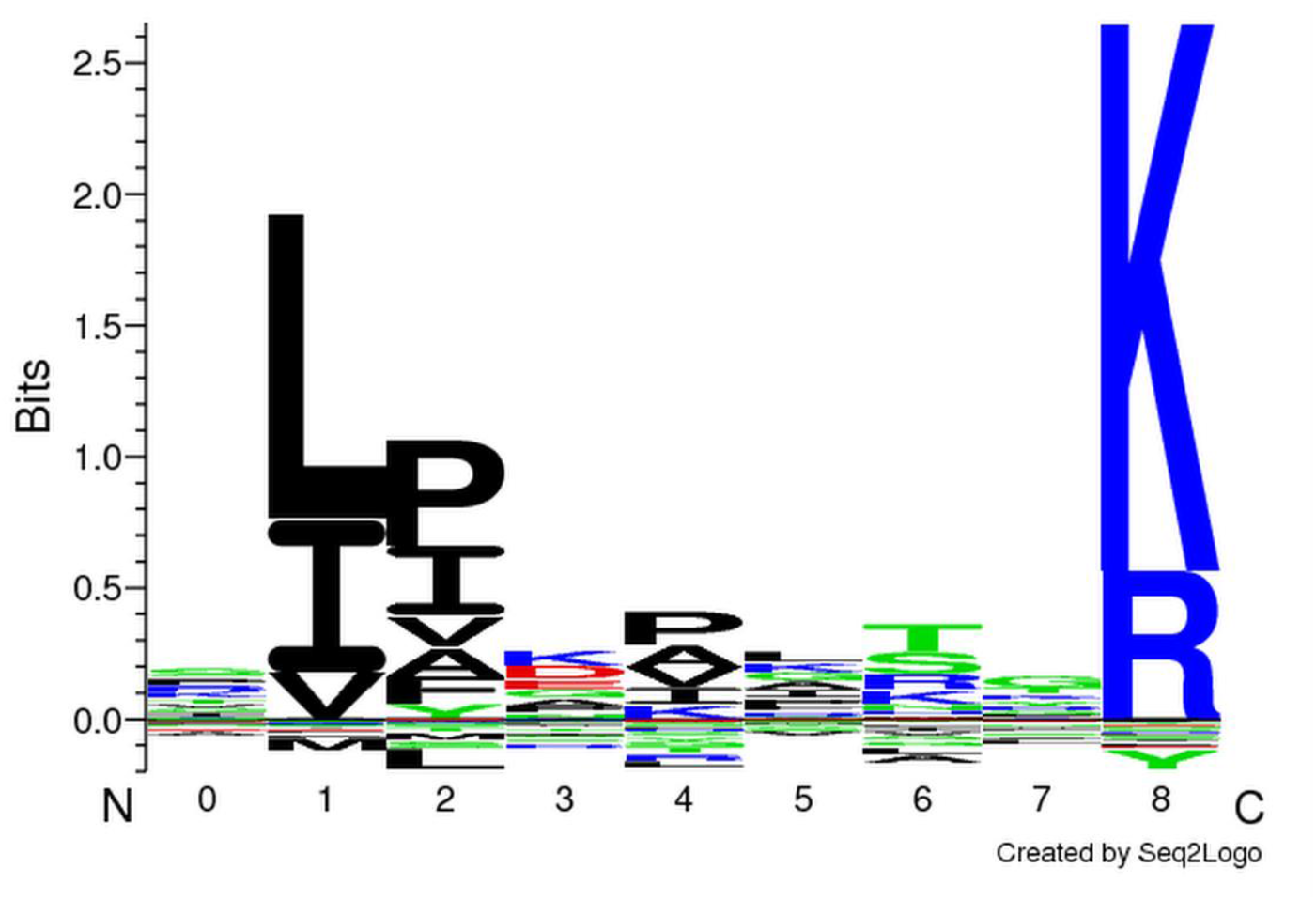
NNAlign binding motif for model trained against INS-1E peptides. The model was trained against RT1.A-bound and random natural peptides.

